# Simulation model for dynamics of three types of annual plants

**DOI:** 10.1101/074393

**Authors:** Andrzej Pękalski

## Abstract

A Monte Carlo type model describing dynamics of three pairs of annual plants living in a homogeneous habitat is presented and discussed. Each plant follows its own history with growing, fecundity and survival chances determined individually as functions of the plant’s condition and environment. The three plants - *Valerianella locusta, Mysotis ramosissima* and *Cerastium semidecandrum* differ by the weight of their seeds, which in the model determines the competition preference. Heavier seeds have a better chance for germination from a site containing seeds of different plants. Better colonisers produce more seeds and disperse them over a larger distance. I show that without absolute asymmetry in the impact effects between better competitors and better colonisers and in a spatially and temporarily homogeneous habitat, coexistence of species is possible, however only in a limited time. This is different from statements coming from models using mean-field type methods. I demonstrate also that in a system of two species clustering of plants of the same type are more frequent. From the calculated survival chances of seedlings and adult plants it follows that elimination of plants occur mostly at the early stages of the plants life cycle, which agrees with the field data.I show that this competition/colonisation trade-off model is sufficient to maintain coexistence and I determine the conditions for dominance of one type of plants. I show that the time of extinction of the weaker species goes down with increasing observation time as a power function with the exponent independent of the type of plants.

## 1 Introduction

It has been remarked by Crawley (1990) that description of plants dynamics is not an easy task. One of the central issues is the problem of coexistence of species, important from practical and challenging from theoretical point of view. Several factors promoting biodiversity have been proposed -spatial heterogeneity (Yu and Wilson 2001; Turnbull et al. 2004), tolerance-fecundity trade-off (Muller-Landau 2010) or disturbances (Roxburgh et al. 2004; Miller et al. 2012; Seifan et al. 2012). Another possibility is the competition/colonization trade-off developed by many authors (see e.g. (Levins and Culver 1971; Tilman 1994; Holmes and Wilson 1998)). The idea is that species which are inferior competitors can coexist with superior competitors if they are better colonisers, i.e. have a higher dispersal rate (more seeds distributed over a larger distance). The question is - what makes a plant a better coloniser or a better competitor. It has been speculated by Rees (1995); Turnbull et al. (1999; 2004) that seed weight could be such a factor.

Coomes et al. (2002) used the neighbourhood models introduced earlier by Pacala and Silander Jr (1985; 1990) for the dynamics of two annual plants *Aira, praecox* and *Erodium cientarium.* The authors concluded that larger seeded *Erodium* was competitively superior to smaller seeded *Aira.*

The competition/colonisation trade-off problem has been modelled in many ways. Rees et al. (1996) used field data and semi-empirical formulae to predict population abundance for some assumed forms of interactions. Holmes and Wilson (1998) constructed a cellular automata model in which two plants disperse their seeds at different distances. The authors constructed phase diagrams in the plane of the colonisation rates of the two plants. Later Yu and Wilson (2001) investigated the competition/colonization trade-off using a model based on averaged characteristics. They have shown that within their model the competition/colonisation trade-off can maintain biodiversity only in a spatially heterogeneous environment.

In the papers dealing with competition/colonisation trade-off it is generally assumed that better competitors have an impact on worse ones, but not vice-versa and this absolute asymmetry is often considered a necessary condition for explaining coexistence of species (Adler and Mosquera 2000). Likewise, it is often assumed (Wilson and Nisbet 1997) that the better coloniser is spreading its seeds over distances greatly exceeding those of the poor coloniser. Turnbull et al. (2004); Miller et al. (2012) showed that also smaller plants have some influence on the larger ones, and the problem of estimating the distance over which seeds are dispersed is difficult (Nathan and Muller-Landau 2000). It is therefore important to check whether without absolute asymmetry and within the pre-emptive model, the competition/colonisation trade-off could be a sufficient mechanism to maintain coexistence of species in a habitat which is homogeneous both in space and in time (no disturbances nor patches with different living conditions). To the best of my knowledge such a model has never been reported. The predictions obtained with this model are consistent with empirical data and enhance our understanding of the mechanisms acting in plant communities.

In this paper I study the dynamics of a system of three annual plants with different seed weights. My model uses a Monte Carlo simulation technique and has been introduced earlier (Kącki and Pękalski 2011; Pękalski and Szwabiński 2013; Droz and Pękalski 2013). The main features distinguishing it from previous studies of the competition/colonization trade-off mechanisms in annual plants communities is in taking into account all of the features listed below. Each plant is described by functions depending on local characteristics determined by actual condition of the plant and its surrounding. These local conditions, not assumed global functions, determine the chance the plant has for survival and its fecundity. I allow also weaker competitors to have some impact on the better ones and finally I take the seed weight as the factor determining which plant is a better competitor. Incorporating all these features makes my model more realistic and also more complex than the ones using global variables, as it is sensitive to local changes in plants’ characteristics it treats individually each plant. As remarked by Watkinson (1990), dynamical behaviour of plants’ populations in nature depends not only on density-dependent factors but also on a variety of other, density-independent ones, biotic and abiotic. These factors, random by nature, are incorporated into the stochastic features of Monte Carlo-type simulations. Such individual treatment of each plant, despite global homogeneity of external conditions, leads in my model to different local ones, which, as stressed by Crawley (1990) should be regarded as something of even greater importance than temporal variation.

The main object of this study is to show that without strict asymmetry in the competitor-coloniser abilities, coexistence is possible. However apart from that, the model allows to obtain several interesting results concerning details of the plants’ dynamics and the way a weaker species is eliminated.

## 2 Background

I consider a similar system as the one studied by Rees et al. (1996), who used in their model difference equations to describe dynamics of the system and the parameters of the model were rather complex functions indirectly only linked with the observed characteristics of the plants. Out of the four species considered in Rees et al. (1996) differing in the weight of their seeds I have chosen the following: *Cerastium semidecandrum, Myosotis ramosissima* and *Volerianella locusta.* The weight of their seeds are 0.08 mg for *Cerastium,* 0.17 mg for *Mysotis* and 0.80 mg for *Valerianella.*

The size, or weight, of a seed is related to the number of seeds produced by a plant. In Rees et al. (1996) the average number of seeds produced by one plant is given as - 20.3 for *Cerastium* and 7.5 for *Myosotis.* It has been argued by Turnbull et al. (1999) that there exists a simple relation between seed mass and the number of seeds a plant produces

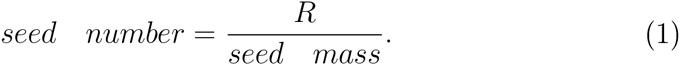
Using *Cerastium* to determine *R* in eq.(l) one can get for the three plants the average number of seeds a plant produces - 20 for *Cerastium,* 9 for *Myosotis* and 2 for *Volerianella.* Rees et al. (1996) showed that the percentage of seedlings survival increases with the seeds size, which indicates that larger seeds have a better chance to germinate, which means that larger-seeded plants are better competitors. Similar statements could be found in other papers (see e.g. Rees (1995); Turnbull et al. (1999; 2004)). The three above mentioned plants, have the same basic requirements for resources (Grubb 1977).

To coexists in a community of several plants those having smaller seeds must compensate this handicap by either their bigger number and/or larger dispersal distance. Although intuitively appealing, the idea that smaller seeds are distributed farther, is not easy to verify experimentally, as described by Nathan and Muller-Landau (2000). Turnbull et al. (1999) noticed that heavy seeds are dispersed over a smaller distance than the light ones. Dispersal distance has an effect on the ability of an inferior species to coexist, as remarked in Holmes and Wilson (1998). It is not clear whether a large difference between the dispersal distance of two competitors fundamentally changes coexistence criteria, or lengthens the time to reach equilibrium (Holmes and Wilson 1998).

It has been shown, see Watkinson and Harper (1978) and references therein, that mortality of plants happens in different periods of the plants’ life cycle, depending on the average number of seeds produced by the plant. When there are just a few of them as for *Vulpia fasciculata,* then the elimination happens at the late stages (flowering) of plant life. This means that the survivorship curve is negatively skewed (Deevey type I) (Deevey 1947). Plants producing many seeds have survivorship curves positively skewed (Deevey type III), meaning that mortality of juvenile plants is high.

Rees et al. (1996) do not give details about the life cycle of the plants investigated by them, but such information could be found in Watkinson and Harper (1978) for *Vulpia fasciculata,* another of the sand dune annuals. It goes as follows:

germination - September - October (about 8 weeks),
growth - November - May (about 28 weeks),
flowering - May - July (about 10 weeks),
After flowering plants are immediately dying. Total lifespan is then about 46 weeks.

## 3 Simplifications

I assume that all my plants - *Cerastium semidecandrum, Mysotis ramosis-sima* and *Valerianella locusta,* have the same demand for one external resource (water) which is divided symmetrically (see below) and their lifespan is 46 weeks, divided into three stages - germination (1 week), growth (44 weeks), flowering and seed dispersal (1 week).

Maximum numbers of seeds produced by a plant, *µ*, are taken as: 25 for *Cerastium,* 10 for *Myosotis* and 5 for *Valerianella.* Only the last one is considered a plant producing such a small number of seeds that plants elimination follows the Deevey I curve. For *Valerianella* time *t*_*x*_ = 22 weeks determines the start of the elimination process, while for the remaining two types of plants it sets the end of it.

Larger seeds have a batter chance to germinate from the same site than smaller ones (Rees et al. 1996; Turnbull et al. 1999). To account for that, the probabilities of choosing a larger seed from a site on which there are seeds from different types of plants are my control parameters. These probabilities could be in a straightforward way linked with competition coefficients of second (logistic) type introduced by Rees and Westoby (1997). Seeds of species with largest seeds, *Valerianella* could remain active for 2 years, for *Mysotis* 3 years and the smallest seeds of *Cerastium* for 4 years. Particular values are not very important, however letting smaller seeds last longer reduces the preference for choosing heavier seeds and decreases the coexistence zone.

## 4 Model

To investigate fully the parameter space their number must be kept low. Therefore I fix the values of most of the parameters of my model at values taken either from the paper by Rees et al. (1996) or Watkinson and Harper (1978). However at the end of the Results section a short estimation of the robustness of the obtained results is given.

The habitat is a square lattice of linear dimensions *L* × *L* with *L* = 100. Each site (plaquette) could be either empty or contain one plant but an arbitrary number of seeds. Time is divided into two units - smaller ones (weeks, denoted by *t*) and larger ones (years, denoted by *T*). All external conditions are reduced to one resource *w*(*t*), which is homogeneous over the whole lattice but vary in time (weeks). All plants have the same demand, *w*_*d*_, for the resource, which is equal to average supply, *w*_0_, taken as 0.5.

The life cycle of plants is composed of three stages - seeds, seedlings and adult plants. Transitions between stages, like e.g. germination of a seed, are determined by the appropriate probabilities depending on the plant fitness. During their life cycle plants’ demands for the resource are compared with its availability. The larger is the difference, the greater is the chance that the plant would be eliminated or would produce less seeds. At the end of their life cycle plants disperse their seeds over areas depending on the mass of the seeds and they die, Choice of a seed for germination depends on the number of seeds on a given plaquette and the preference of choosing a seed for germination. Chosen seed is then put into a germination test, also depending on the fulfilment of the demand for the resource. If the seed passed the test it becomes a seedling and in the next week it turns into an adult plant, which lives during 44 weeks. Therefore resource availability determines all steps of the plant’s life cycle.

At the beginning of simulation 1000 plants of each of the two types, are put randomly on an empty lattice. Another possibility of studying coexistence has been proposed by Miller et al. (2012), when a small number of intruders is put inside an existing population.

Plants compete with their nearest neighbours within the von Neumann neighbourhood (henceforth denoted by NN) for the resource in a symmetric way (Weiner 1990), i.e. the supply on the central plaquette, where the plant grows, is divided by the number of plants in NN. On a square lattice, which I am using, there are 4 such NN. Small variations in time of the external resource are described as

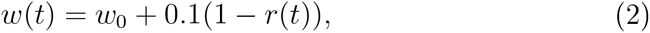
where *w*_0_ is the average supply, *r* ∈ [0,1] here and afterwards is a random number taken from a uniform distribution. The factor 0.1 is chosen small to reduce the chance for major perturbations, which would require separate studies. The amount of the resource a plant *i* with *nn*_*i*_ nearest neighbours may get in a week *t* is

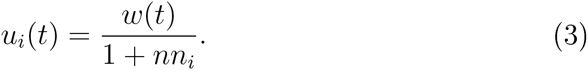
This is compared with the plant’s demand *w*_*d*_

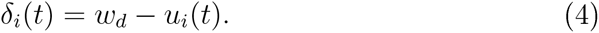
*δ*_*i*_(*t*) is then used to calculate the chance the plant has to survive (if this happens in the elimination period, i.e. first weeks for *Cerastium* and *Mysotis* and last weeks for *Valerianella*)

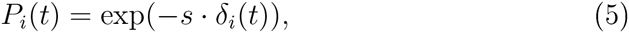
where *s* is the selection pressure, taken in the simulations as 0.05. If a new random number *r* ∈ [0,1] ≥ *P*_*i*_, the plant is eliminated and another plant is randomly chosen for inspection. The calculated value of *P*_*i*_ is used also to estimate the plant’s fecundity, *f*_*i*_,

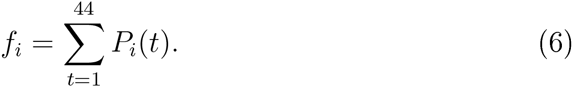
The summation is taken over all weeks of plants’ adult life. The calculated value of the fecundity determines the number of seeds the plant produces

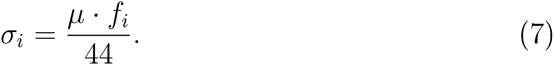
Here *µ* is the maximum number of seeds that a plant of a given type could produce (Seifan et al. 2012). Using the same function *P*_*i*_(*t*) for the plant’s survival chance and the number of seeds it produces is justified, since as shown in Shirley (1929) the number of seeds depends on how well the plants’ demands for resources have been met at all stages of its life cycle. This procedure of random choosing plants and determination of their survival and fecundity is repeated as many times as there are plants in the system. At the end of a year all surviving plants disperse their seeds in the numbers calculated from eq.(7) over distances typical for their type - in the four NN and the plaquette on which the plant grows for *Valerianella,* in 8 + 1 sites for *Mysotis* and in 24 + 1 plaquettes for *Cerastium.* After dispersing the seeds plants die. A sweep over all lattice is done and all seeds older than the predefined value typical for the plant type are removed. This corresponds to different time limits of the respective seed banks. Next year starts with the germination phase. In it all plaquettes containing seeds are randomly visited and one seed for germination is chosen. In the case of one type of plants the choice is random. When there are two types, the lottery model of Chesson and Warner (1981) is applied. If there is *s*_*V*_ seeds of plant V and *s*_*M*_ seeds of type M, then the probability of choosing a seed of type V is

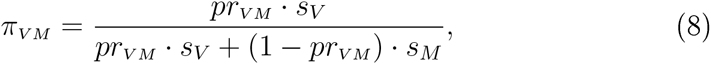
where the parameter *pr*_*VM*_ is a weight describing preference in choosing a seed of type V from a pair (V, M). Analogous formula, *mutatis mutandis,* is used for other pairs of plants. The selected seed is then put to the germination test - calculation of *P*_*i*_ from eq. (5) and comparing it with a random number *r*_*i*_ ∈ [0,1]. If *r* ≤ *P*_*i*_ the seed becomes a seedling and no further germination from this plaquette is possible in this year. When all plaquettes containing seeds have been visited, the germination phase is over and all seedlings become adults plants.

My control parameters are the competition powers of the plants - the weights *pr*_*kl*_ in the probabilities *π*_*kl*_ of choosing a larger seed when on the same plaquette seeds of different types are present. Inb my model the effect of preference is increasing with the density of plants in agreement with (Rees 1995; Turnbull et al. 1999), since at low densities the chance that a plaquette will contain two types of seeds is small.

The main, and very important, difference between my model and the one used in Rees et al. (1996) is that they described the dynamics of the plants via a set of difference equations using globally defined parameters. I am investigating here many aspect (out-performance of one species, zones of coexistence, elimination time, seedlings vs. seeds success) not touched upon in Rees et al. (1996).

## 5 Results

Dynamics of one type of plants growing alone is quite similar for the three species. After an initial stage, which is shortest for the best coloniser and longest for the best competitor, a stationary state is reached. I investigate a pairwise system composed of two annuals - either *Valerianella* and *Mysotis* (V,M), or *Valerianella* and *Cerastium* (V,C) or *Mysotis* and *Cerastium* (M,C). In the first case the control parameter is the probability, *pr*_*VM*_., in the second case it is *pr*_*VC*_ and in the third case *pr*_*MC*_.

Plots showing temporal dependence of the plants’ abundance were obtained after averaging over 25 realisations, i.e. 25 different spatial distribution of the same initial population. The simulations were run for 60 years. In some cases, like for the pair (V,C) and *pr*_*VC*_ = 0.90, it seems that a stationary state is reached. However at times of the order of hundreds of years one species, (C), eliminates the other (V). This agrees with earlier observations (Chesson and Huntly 1997). Hence, what I present here is a transient stage, which could last for several decades. How the time at which the data are collected influences the results, is discussed below. Sets of parameters values which well illustrate the tendencies and the role of the parameters are chosen.

Figure 1, shows the time (in years) dependence of the abundance of plants for the pair (V,M) and for three values of the weight *pr_VM_*. Figures 2 and 3 show the same for the pairs (V,C) and (M,C). The features in all cases are similar and with increasing the probability of choosing a seed of a poorer coloniser its number is growing and if the probability is high enough the plant could be the dominant one. At earlier stages we observe a negative relation between seed size and abundance, as found by Rees (1995). At the beginning in the habitat there are only initially put plants and a better coloniser is winning, regardless of the preference in seed selection, since the total density of plants is low and plaquettes containing both types of seeds are rare. With increasing density the situation changes and the competition advantage is more important. Figure 4 shows spatial organisation of plants at the end of simulations (60 years) for values of the weights at which the species coexist. To see better the locations of plants, only a fragment of the lattice is shown. Some clustering of plants of the same type is observed, also for *Cerastium* which disperses its seeds rather far.

**Figure 1.**
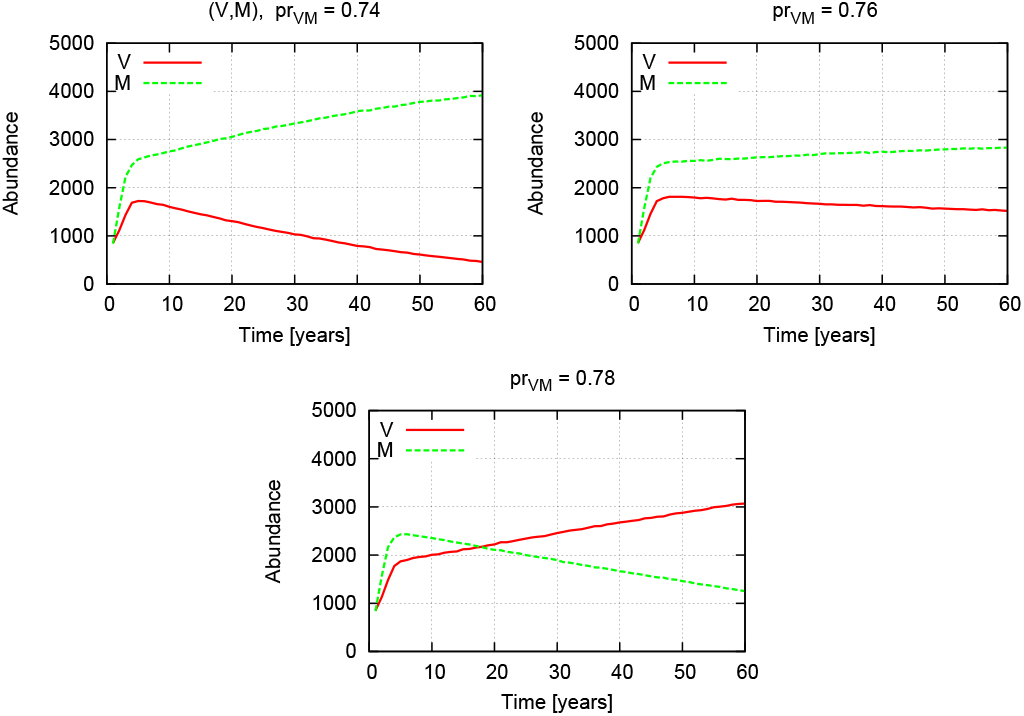
Temporal dependence of the abundance of plants *Valerianella* and *Mysotis* for three values of *pr*_*VM*_ - preference of choosing a *Valerianella* seed from a site containing both types of seeds. See eq.(8).

**Figure 2.**
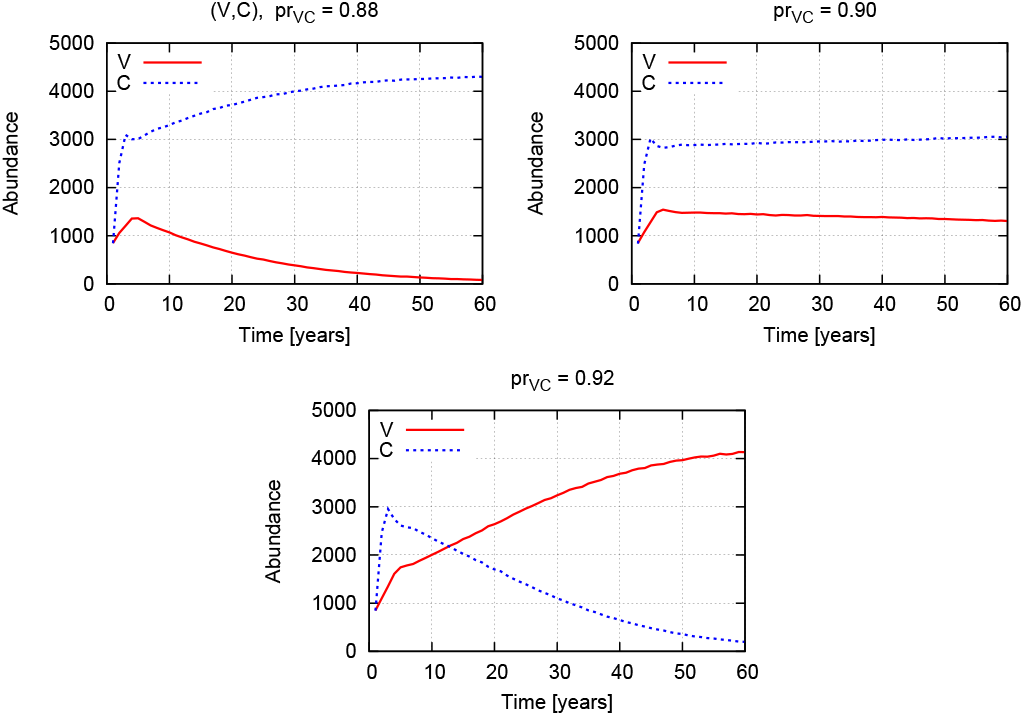
Same as in Figure 1, but for the pair (V,C).

**Figure 3.**
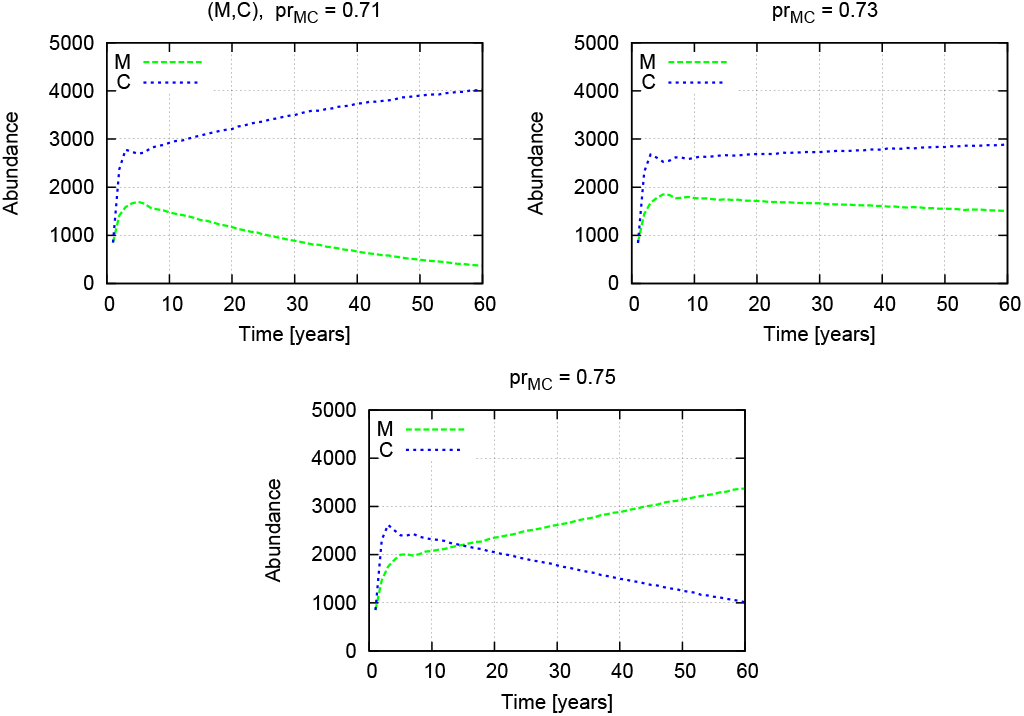
Same as in Figure 1, but for the pair (M,C).

**Figure 4.**
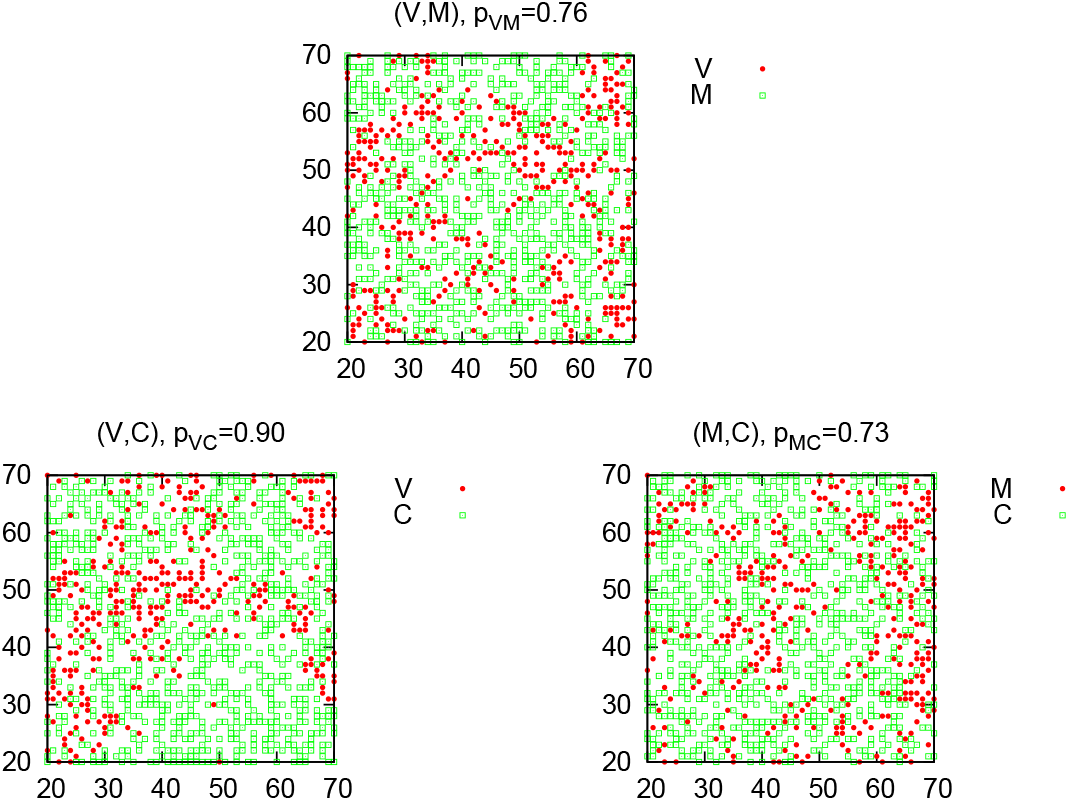
Spatial arrangement of plants at the end of simulations (60 years) for three pairs of plants and values of the *pr*_*kl*_ parameters corresponding to the coexistence region. A part of the total lattice is shown.

Clustering of plants of the same species could be investigated in more detail by calculating the average fraction of plants in NN of the same type as the central plant, compared to plants of both types for the same central plant. It is evident that with increasing the percentage of plants of a given type also the fraction of alike plants in the nearest neighbourhood will grow. To compensate for this effect I introduce a function *ϱk* (k = V,M,C) defined as

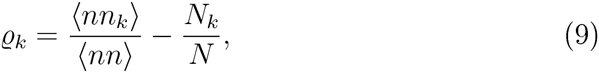
where ⟨*nn*_*k*_⟩ is the average number of nearest neighbours in NN of the same type as the central plant *k,* ⟨*nn*⟩ is the average number of both types for the same plant, *N*_*k*_ is the abundance of plants of type *k* and *N* is the total number of plants. When *ϱk >* 0 intra-species interactions prevail and are more frequent than following from simple density effect.

Figure 5 shows it for the three cases. The general features are in all cases quite similar. Strong preference for clustering of plants of the same type is observed in the coexistence zone. There are therefore more intra-specific interactions than inter-specific ones, as remarked in Rees et al. (1996). When one of the species nearly or completely eliminated the other one, *ϱk* approaches zero, as seen from eq.(9).

**Figure 5.**
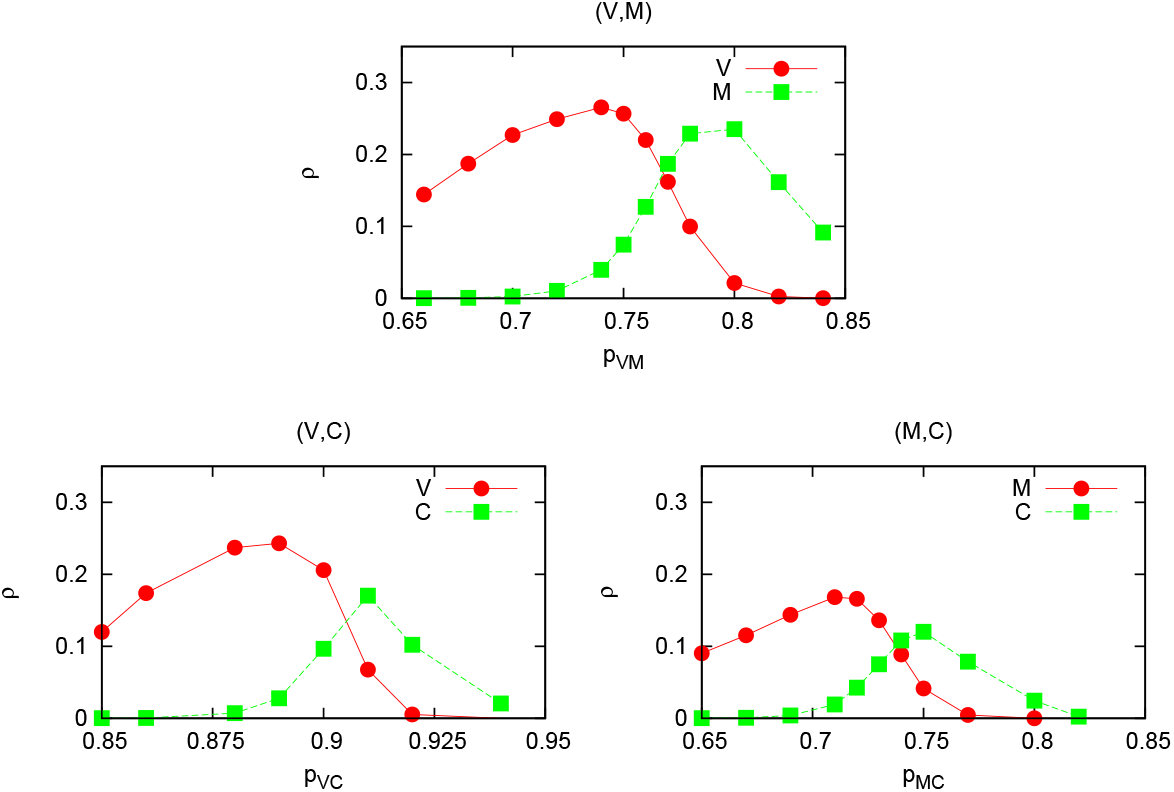
Reduced fraction of the same plants in the von Neumann neighbourhood (see eq.(9)) taken after 60 years, for the three pairs of plants and as functions of *pr_kl_.*

It is interesting to check at what stage - adult plants or seedlings the plants are more often eliminated. To this end I have calculated the seeds (*SN*) and seedlings (*SK*) successes, defined as follows. The seeds success is the ratio of the number of successful germinations to the number of plaquettes with seeds. From a plaquette only one seed can germinate in a given year, hence it is the number of plaquettes with seeds which is essential, not the number of seeds. The seedlings success is the number of adult plants which survived till the end of simulations and produced seeds, divided by the number of seedlings. Figure 6 shows the successes as functions of the preferences in choosing larger seeds and for the three pairs of plants. From Figure 6 it follows that the seed success, *SN* (empty symbols), depends heavily on the value of the weight parameters, while the seedlings success, *SK,* remains much higher and its dependence on the parameters is weaker. The obtained results (seedlings survival chance for all considered here cases about 75 %) agree quite well with the survival estimates given in Watkinson and Harper (1978) for *Vulpia fasciculata* - about 70 %. The overall conclusion is that the abundance of plants is controlled at the seed level, not by killing adult plants. It confirms earlier statements (Watkinson 1980; Levine and Rees 2002).

**Figure 6.**
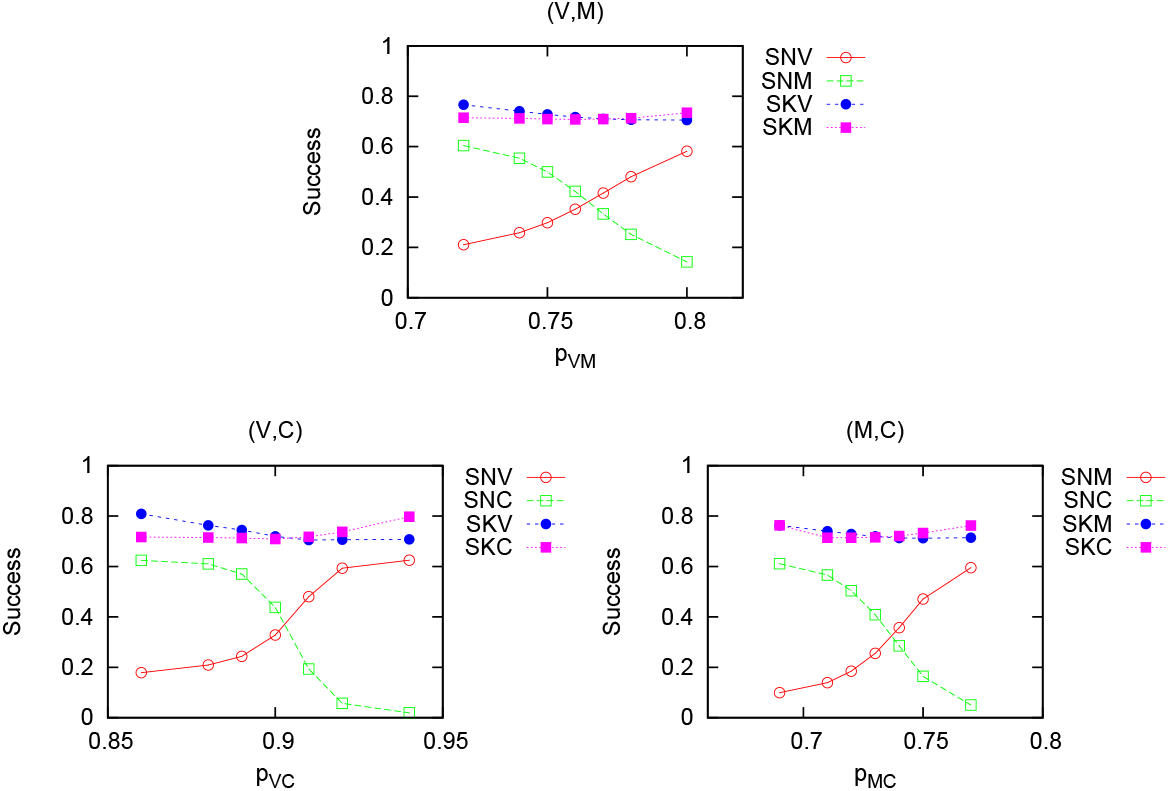
Seeds (open symbols) and seedlings successes as functions of the *pr*_*kl*_ parameters.

The results presented so far were obtained for simulations lasting 60 years. Since however the system is not in a stationary state, it is necessary to check how the results change when the observation time is different. The information could be useful to determine trends present in a real habitat.

The dependence of the number of plants on the time of collecting data is shown in Figures 7, 8 and 9 for the three pairwise ‘experiments’. The coexistence region shrinks with time, meaning that coexistence is possible only for a more and more restricted range of the weight parameters. A question arises whether the rate of disappearance of the coexistence regime is the same for all considered systems, or it is specific.

**Figure 7.**
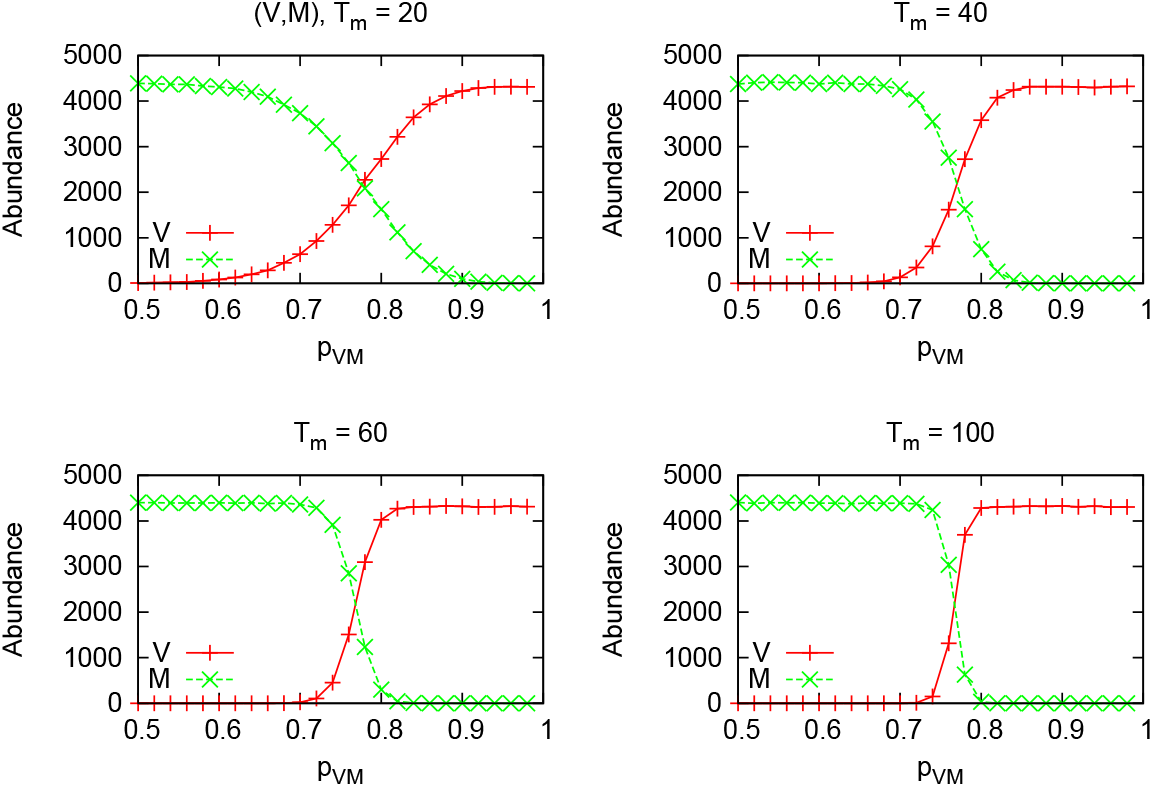
Abundance of plants taken at different simulation times, *T*_*m*_, for the pair (V,M), as a function of *pr*_*VM*_.

**Figure 8.**
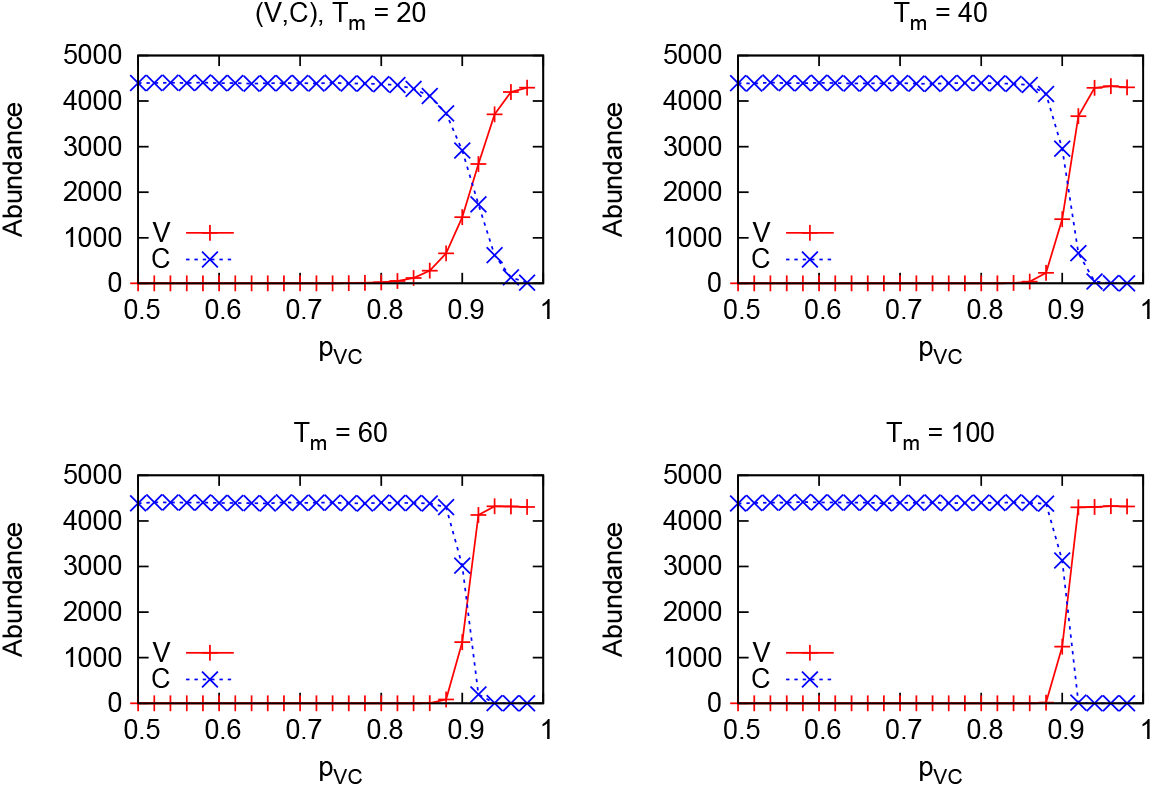
Same as in Figure 7, but for the pair (V,C).

**Figure 9.**
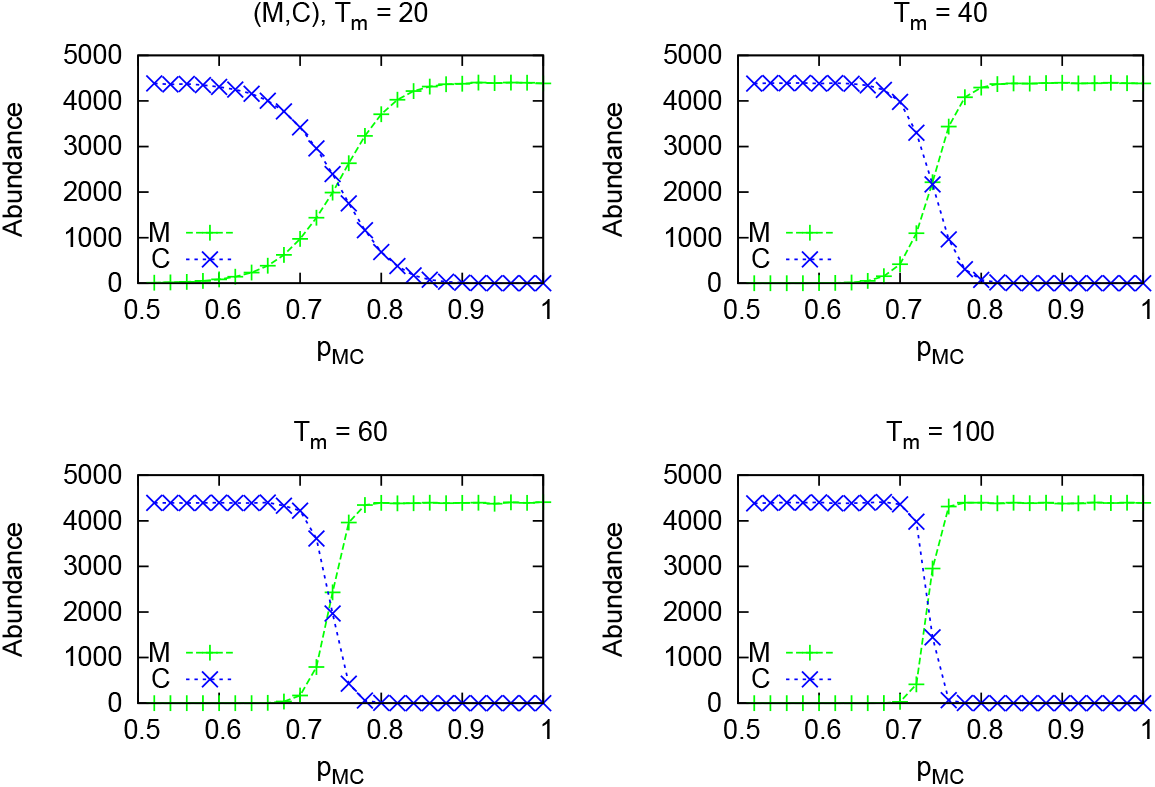
Same as in Figure 7, but for the pair (M,C).

Figure 10 shows the width, Δ, of the coexistence range as a function of the observation time, *T*_*m*_, in years, on a doubly logarithmic plot. Apart from a pre-factor, all three cases follow a power dependence with the same exponent

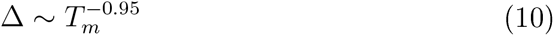
Hence, shrinking of the coexistence region does not depend on the details of the plants characteristics and has a universal character. Finally, Figure 11 shows the coexistence regime dependence on the parameters *pr*_*kl*_ (k,l = V,M,C) and time of observation, *T_m_.* All three cases have the same qualitative character and coexistence in all is possible. When a poor coloniser (*V*) competes with a very good one (*C*), only when the preference of choosing a *V* seed is extremely large, the *V* can eliminate *C.*

**Figure 10.**
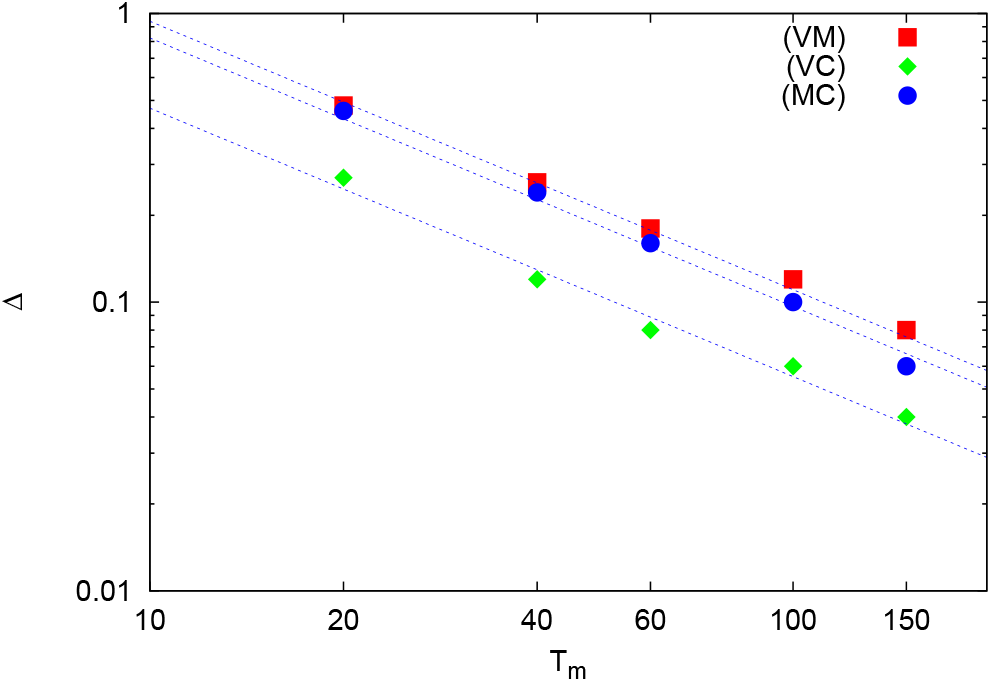
Width, Δ, of the region of the parameters *pr*_*kl*_ values where coexistence is possible, as a function of the observation time, *T*_*m*_. Double logarithmic scale.

**Figure 11.**
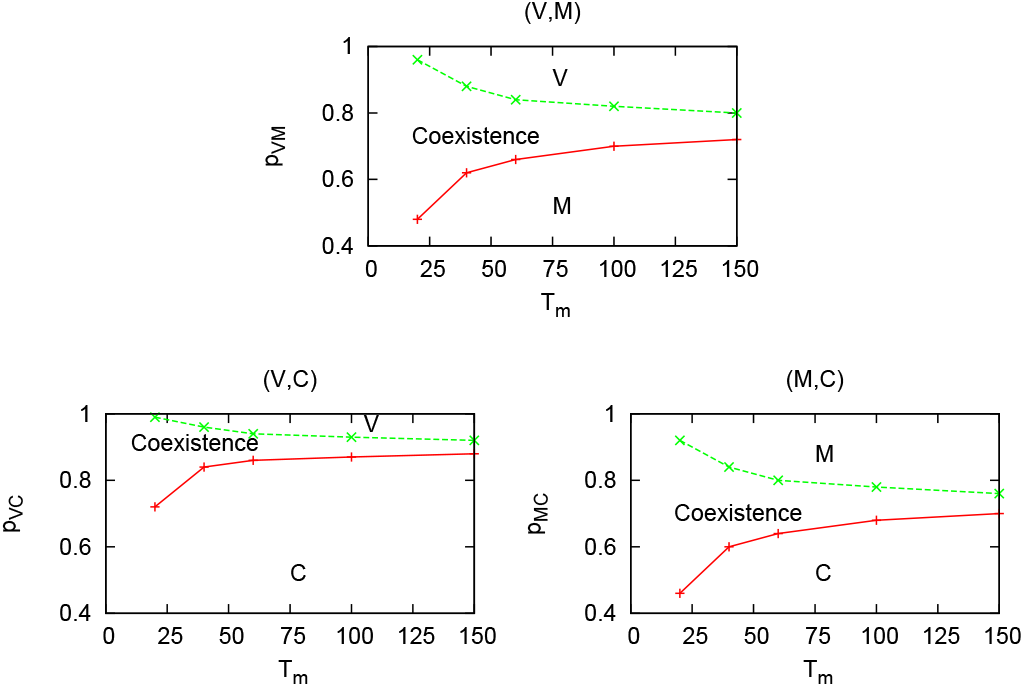
Phase diagram showing in the (*pr*_*kl*_, *T*_*m*_) plane, the coexistence regime and regions where only one type of plants is alive.

### 5.1 Robustness

I discuss here effects of some changes of the parameters of the model, which were not shown in the figures. Producing plots for all values of the control parameters would be not practical, since e.g. for all cases of small *pr_VC_ Valerianella* would be eliminated, because in that range it is both a worse competitor and a worse coloniser. Therefore only significant ranges of the parameters, well illustrating the tendencies, and the role of the parameters, have been chosen

I have verified that populations from different realisations (different initial distribution of plants) have almost identical history Hence, the dynamics is robust with respect to changes of initial conditions and the plots show typical behaviour. Increasing the value of the selection pressure from the adopted here *s* = 0.05, leads to very fast extinction of plants, while making it smaller greatly reduces influence of the environment. The size of the lattice (10 000 plaquettes) is sufficient to eliminate all stochastic extinctions and provides reasonable execution time of the simulations. Keeping the ratios of the maximum number of seeds a plant of a given species could produce (here 5, 10, 20 for the *Valerianella, Mysotis* and *Cerastium,* respectively) fixed and changing the numbers, like to (10,20, 40) does not influence the results in any way.

The area over which seeds are distributed is one of the parameters of the model which is difficult to verify experimentally. Therefore, apart from the reported above case when the areas were 5, 9 and 25 plaquettes for *Valerianella, Mysotis, Cerastium*, respectively, I have run simulations for the following areas - [5, 9, 13] and [9, 13. 25] for the three plants. In all cases coexistence has been found in the regions quite similar to the ones for the areas [5, 9, 25]. Therefore the particular values of the distances over which the seeds are distributed, as long as the sequence is preserved, are not crucial for establishing coexistence.

Robustness of the results presented above with respect to changing plants features indicate that even when the plants are described by more complex characteristics, coexistence is still possible.

## 6 Conclusions

For coexistence to exist it is often assumed that either complete assymetry between a good coloniser and good competitor was necessary (Turnbull et al. 1999; Yu and Wilson 2001), or a better coloniser must distribute its seeds much farther (Tilman 1994; Wilson and Nisbet 1997).

A simulation model of plants dynamics developed earlier (Kącki and Pękalski 2011; Droz and Pękalski 2013) is applied here to study systems of three annual plants - *Valerianella, Myosotis, Cerastium* living in pairs. The habitat is spatially homogeneous, hence coexistence could not appear via patch preference. Also the external conditions remain unchanged in time, therefore diversity of species could not be linked to disturbances. A plaquette on which grows a plant could not be invaded by another one. The mechanism responsible in my model for coexistence is the competition/colonisation trade-off, with seed size as the factor differentiating between a good coloniser and a good competitor. I proposed a simple definition of a better competitor as a plant which has a heavier seeds than the other plant and such seeds are more often chosen from sites containing both types of seeds. This is based on earlier observations (Rees 1995; Rees et al. 1996). A better coloniser produces more seeds and disperses them farther. Although in my model the habitat is homogeneous, local conditions differ from plant to plant, as differ the number of nearest neighbours interacting with the central plant.

The obtained results indicate that at the early stages of colonisation of an empty habitat, the best coloniser always is the dominant species, as remarked in Rees (1995), regardless of the value of the parameters. In an intermediary period (5–15 years) many patterns are possible and the final state depends on the values of the parameters.

I have shown that both differences (asymmetry and spreading distance) could be much smaller to allow for coexistence of species and that the competition/colonisation trade-off is a sufficient mechanism to establish the coexistence of two species, albeit for a restricted time interval.

This difference in the conclusions about the role of the competition/colonisation trade-off in earlier papers and this one could be due to the fact that in mean-field type models all conditions are global, hence all functions determining the dynamics of plants and their characteristics are also global and local effects are lost. In investigations of any complex system, biological or else, fluctuations of the basics quantities are very important. They are, by construction, totally neglected in a mean-field type approach, which in some cases could give therefore even qualitatively wrong results. For example in the mean-field model unfavourable conditions apply to all plants and the population may go extinct. In a simulation model, like this one, and in nature, conditions are determined locally and they affect only a small group of plants not endangering the whole population. Only inclusion of individual treatment of objects, updating their characteristics according to local conditions and allowing for some stochasticity can reasonably well mimic the nature. The importance of agent based type of modelling has been discussed e.g. in Grimm et al. (2005).

I have demonstrated that in my model the coexistence of species is a transitory phenomenon and I have, to the best of my knowledge, for the first time investigated how fast the weaker species disappears. I have shown that it goes as a power-type function and that the effect does not depend on the species characteristics, hence it is universal.

In my model the abundance of plants is controlled at the seeds level, while killing adult plants is rare, as found in Watkinson and Harper (1978); Levine and Rees (2002). Clustering of plants of the same species is observed, specially when the number of plants in a pair is similar, in agreement with earlier observations (Rees et al. 1996). The region of values of the preference parameters over which coexistence is possible, goes down with increasing observation time as a power function with the exponent independent of the plants characteristics.

I have shown that the difference in the dispersal distance between species could change coexistence chance and not only lengthen the time to reach equilibrium, which answers the question made in Holmes and Wilson (1998). Calculated in my model estimations of chances a seed has to germinate, grow and live long enough to produce seeds, agree with the data for *Vulpia fasciculata* (Watkinson 1990). My model predicts also that there are more intra-specific interactions than inter-specific, as conjectured earlier in Rees et al. (1996).

Obtained results are robust against changes of the plants’ characteristics and therefore have a general character. As such they could be used to determine trends in real ecosystems.

It is, at least conceptually, not difficult to include into the presented model other factors neglected here - spatial and/or temporal inhomogeneities, different tolerances of plants to a stress coming from a shortage of the resource, two or more types of the resources or more detailed characteristics of plants (different size, demands etc.).

## Acknowledgement

I am greatly indebted to J. Lukaszewicz for valuable discussions.

